# Cortical structure of neural synchrony and information flow during transition from wakefulness to light non-rapid eye movement sleep

**DOI:** 10.1101/2022.03.09.483562

**Authors:** Joline M. Fan, Kiwamu Kudo, Parul Verma, Kamalini G. Ranasinghe, Hirofumi Morise, Anne M. Findlay, Keith Vossel, Heidi E. Kirsch, Ashish Raj, Andrew D. Krystal, Srikantan S. Nagarajan

## Abstract

Sleep is a highly stereotyped phenomenon, requiring robust spatial and temporal coordination of neural activity. How the brain coordinates neural activity with sleep onset can provide insight into the physiological functions subserved by sleep and pathologic phenomena associated with sleep onset. We quantified whole-brain network changes in synchrony and information flow during the transition from wake to non-rapid eye movement (NREM) sleep using magnetoencephalography imaging in healthy subjects. In addition, we performed computational modeling to infer excitatory and inhibitory properties of local neural activity. The sleep transition was identified to be encoded in spatially and temporally specific patterns of local and long-range neural synchrony. Patterns of information flow revealed that mesial frontal regions receive hierarchically organized inputs from broad cortical regions upon sleep onset. Finally, biophysical neural mass modeling demonstrated spatially heterogeneous properties of cortical excitation-to-inhibition from wake to NREM. Together, these findings reveal whole-brain corticocortical structure in the sleep-wake transition and demonstrate the orchestration of local and long-range, frequency-specific cortical interactions that are fundamental to sleep onset.

## INTRODUCTION

The transition to sleep involves a set of neural processes that reliably promotes a behavioral state-change and sets the stage for essential functions of sleep. Classical lesion and stimulation-based studies have identified the sleep-wake transition as modulated by distributed circuits originating in the brainstem and extending to the hypothalamus, thalamus, and basal forebrain^1,2^. Such bottom-up regulation of sleep and wake has led to a model whereby cortical structures are globally activated by subcortical structures during state transitions. More recent studies, however, have identified the presence of local cortical dynamics during sleep^3–7^ and furthermore suggest that the neocortex may even play a top-down role in sleep-wake regulation^8,9^. To this end, we pose the questions: Is there a structured coordination between local and long-range cortical dynamics during the transition from wake to light NREM? If so, what are putative mechanisms mediating the local cortical dynamics? The elucidation of coordinated local and long-range cortical structure may ultimately shed light on cortically-based physiologic phenomena, such as emotional regulation^10^, memory consolidation^11^, and recovery^12^, and pathologic phenomena of sleep onset, such as state-dependent epileptic spiking or insomnia.

For many decades, sleep electrophysiology has been characterized by recordings from a limited number of scalp electrodes, as traditionally used in polysomnography. Meanwhile, fMRI/PET^13–15^ studies have provided key anatomical insights on whole-brain neural activation patterns of sleep. Such large-scale network analyses primarily using fMRI have revealed widespread increases in corticocortical connectivity and decreases in thalamocortical connectivity during light non-rapid eye movement (NREM)^16–18^; deep NREM has been associated with widespread decreases in functional connectivity (FC) and increased thalamocortical connectivity^16,17^. The diffuse increase in corticocortical FC from wakefulness to light NREM has been hypothesized to lead to reduced information integration, underpinning the loss of consciousness with sleep^19,20^. While fMRI/PET offer high spatial resolution, these imaging modalities indirectly probe neural activity (e.g., fMRI measures blood flow and PET measures neural metabolic rate) and are limited by low temporal resolution.

Complementing prior fMRI methods, several key MEG/EEG^21–23^ studies have elucidated frequency-specific patterns with sleep onset, including the robust increase in low frequency oscillations in midline anterior brain regions, which later propagate posteriorly^22^. The increased corticocortical connectivity with sleep onset has been furthermore observed across multiple frequency bands using band-limited power correlations of MEG/EEG sensor time series^24^. To date, however, MEG/high-density EEG studies have been limited by the lack of a comprehensive assessment of simultaneous local and long-range neuronal synchrony and information flow with transition from wakefulness to sleep. In addition, amidst findings of global increases and decreases in FC, salient long-range cortical interactions have not previously been observed, which may reflect prior methodological limitations including: limited measures of connectivity due to the lack of source reconstruction^24^; temporal blurring due to the lack of simultaneous EEG for sleep-scoring in MEG studies; and correlational connectivity methods that may introduce effects of volume conduction or other sources of spurious connectivity.

In this study, we sought to identify local and long-range cortical structure that underlie global state transitions from wake to light NREM. We pursued three aims: 1) to determine which cortical areas are preferentially activated by local synchronization during sleep onset using MEG imaging; 2) to simultaneously determine what cortical regions facilitate long-range synchronization observed upon the transition to sleep; and 3) to infer mechanisms underlying the observed network physiology using biophysical models. We hypothesized that there are frequency-specific, spatial patterns of corticocortical synchronization and information flow upon transition to NREM. Furthermore, in the context of spatially-specific cortical patterns of neural synchronization, we hypothesized that the underlying corticocortical structure represents spatially heterogeneous excitatory-to-inhibitory activity during light NREM. To test these hypotheses, we leveraged the high spatial-temporal sensitivity of MEG imaging combined with simultaneous surface EEG to quantify spatial-temporal patterns of local and long-range neural synchrony and information flow upon sleep onset in a cohort of healthy individuals.

## RESULTS

### Local neural synchrony across sleep states

Local neural synchrony, as measured by the spectral power, was averaged across all parcellations and revealed an expected alpha peak during wakefulness and a shift toward delta frequencies with transition to N1 and N2 (Fig. 1A). Source reconstructions of the mean normalized spectral power demonstrated increased delta power predominantly over the bilateral prefrontal cortices with transition from wake to NREM (Fig. 1B). The anterior-posterior gradient in the alpha frequency band, as well as the central beta power, attenuated with transition from wake to light NREM (Fig. 2s). The spectral reconstructions are consistent with prior findings^25^ and demonstrate the accurate acquisition of sleep-wake states.

**Figure 1.**
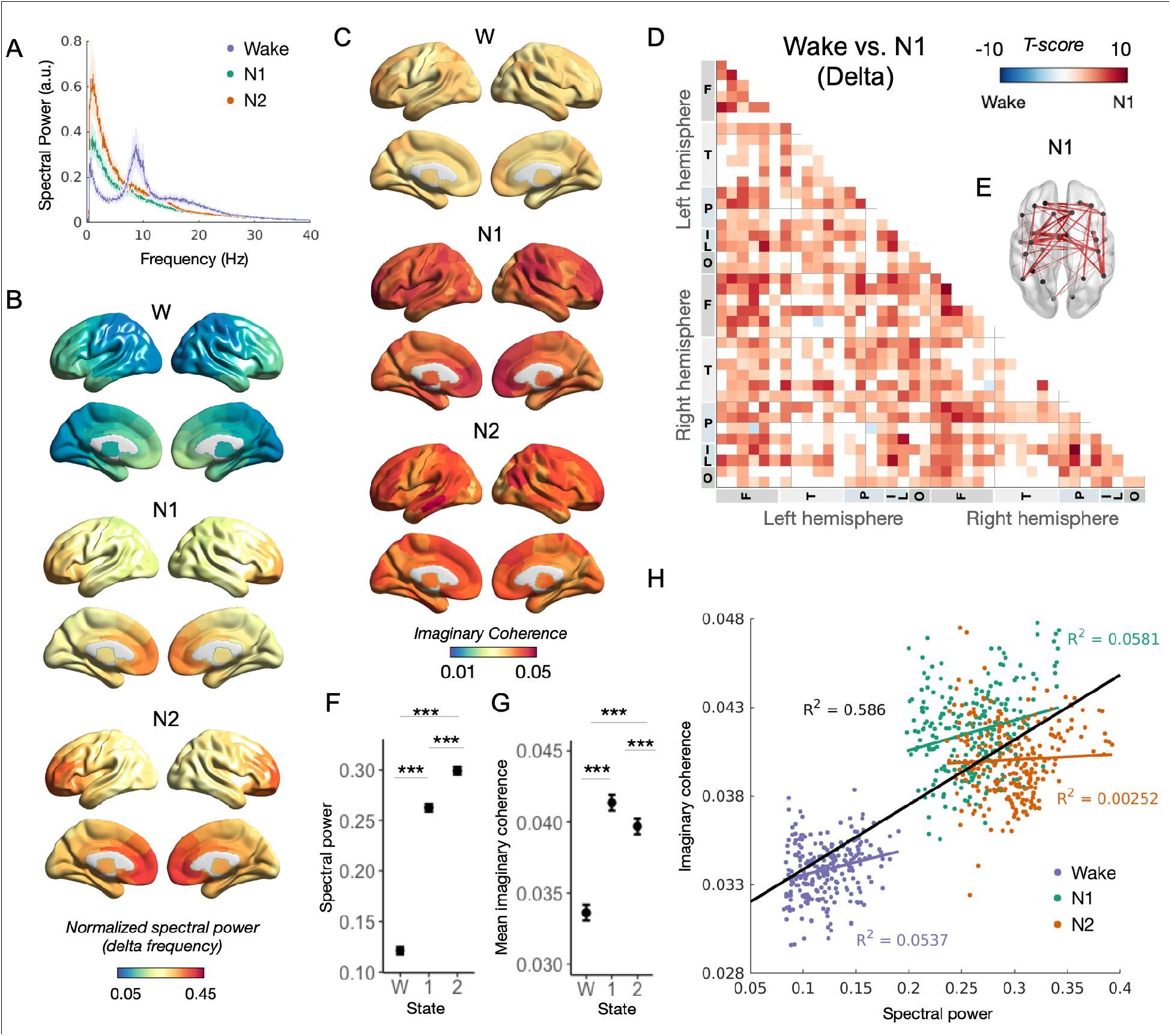
Spatial maps of mean local and long-range synchrony between sleep-wake states in the delta frequency band. A) Mean normalized spectral power, averaged across regions and patients in wake (purple), N1 (green), and N2 (orange). Light shading represents the standard error across patients. B) Spatial maps of mean regional local synchrony (as measured by normalized spectral power) across sleep-wake states. C) Spatial map of mean regional long-range synchrony within the delta band (as measured by imaginary coherence) across sleep-wake states. D) T-score map of differences in long-range synchrony between W and N1 within the delta band. Lobar regions delineated on x- and y- axis are further described in Table 1s. E) Top 50 highest positive functional connections, favoring N1. F) Mean global spectral power of each sleep-wake state in the delta band, averaged across regions and patients. Error bars represent the 95% CI from the LS-mean. G) Mean global long-range synchrony for each sleep-wake state in the delta band, averaged across region and patients. Error bars again denote 95% CI from the LS-mean. H) Association between mean regional long-range synchrony (as measured by imaginary coherence) and mean regional local synchrony (as measured by normalized spectral power) in the delta band. Each point represents an individual ROI within the spatial map. Linear regression and correlations of the mean regional long-range and local synchrony measures are provided for within sleep-wake state and across state (black) populations.

**Figure 2.**
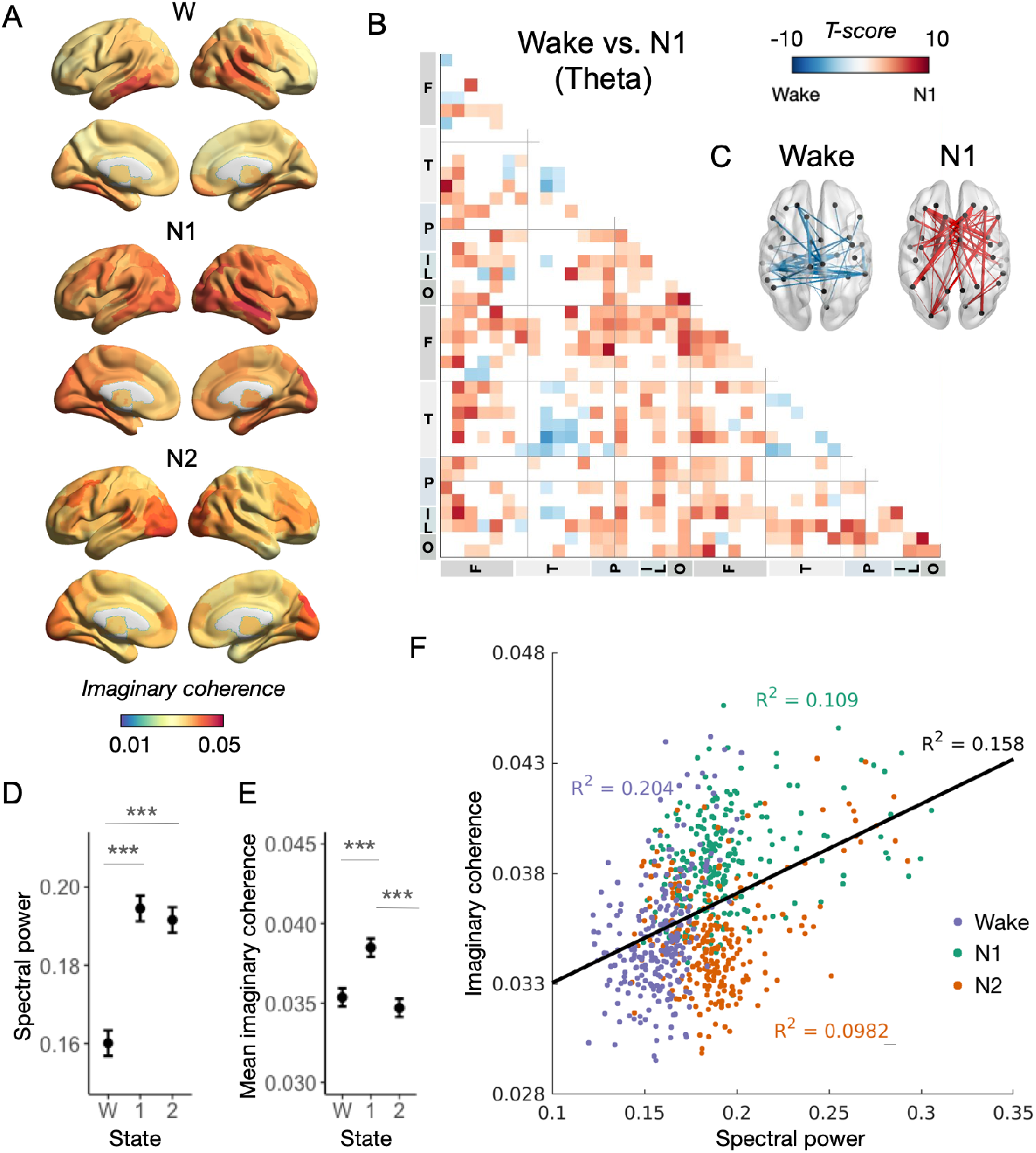
Spatial maps of mean local and long-range synchrony between sleep-wake states in the theta frequency band. A) Spatial map of mean regional long-range synchrony in W, N1, and N2 within the theta band. B) T-score map of differences in regional long-range synchrony between W and N1 within the theta band. C) Top 50 highest positive and negative functional connections, favoring N1 (red) and W (blue), respectively. D) Mean global spectral power of each sleep-wake state in the theta band, averaged across regions and patients. Error bars represent the 95% CI from the LS-mean. E) Mean global long-range synchrony for each sleep-wake state in the theta band, averaged across region and patients. Error bars again denote 95% CI from the LS-mean. F) Association between mean regional local and long-range synchrony in the theta band.

### Long-range neural synchrony maps across sleep states

Mean regional long-range connectivity was mapped onto the Brainnetome atlas to give rise to “spatial maps”, which revealed frequency-specific patterns of mean long-range synchrony of each region in light sleep as compared to wakefulness. Given the prominent shift to delta/theta frequencies with transition from wake to sleep (Fig. 1A), we focused on low frequency long-range synchrony patterns during sleep. Findings for other higher frequency bands are provided in the Supplementary Materials. Within the delta band, mean regional long-range synchrony diffusely increased (Fig. 1C) with prominent long-range connections within and across bilateral frontal regions (i.e. SFG, MFG, PrG) and between frontal-parietal regions (i.e. between MFG, SFG and IPL, SPL; Figs. 1D, E).

Global long-range synchrony, i.e. averaged across all regions, increased from W to N1 (Fig. 1G; t=19.4, p<0.001) and W to N2 (t=15.3, p<0.001), and decreased from N1 to N2 (t=-4.2, p<0.001) in the delta band. The finding of the peak global long-range synchrony in N1 was distinct from that of local synchrony, in which the mean local synchrony monotonically increased from W to N1 and N2 (Fig. 1F). In addition, the relationship between mean regional local and long-range synchrony revealed low correlation within N1 and W states and no correlation within the N2 state (Fig. 1H; purple, green, and red lines; W, R^2^ = 0.054, p<0.001; N1, R^2^ = 0.058, p<0.001; N2, R^2^ = 0.003, p=0.433, respectively). On the other hand, there was a relatively high correlation *across* sleep-wakes states between regional local and long-range synchrony (Fig. 1H; black line, R^2^ = 0.586, p<0.001). These findings suggest that the spatial maps of local synchrony do not strongly reflect that of regional long-range synchrony within classical stages of light NREM; however, mean regional local and long-range synchrony tracts across state changes.

Within the theta frequency band, bilateral frontal and occipital regions (i.e. IFG, MFG, ACC, and MVoCC, LOcC, and Pcun) exhibited increased connectivity, while bilateral temporal regions revealed reduced intra- and interhemispheric connectivity (i.e. STG, ITG, MTG, Hipp and Amy) in N1 as compared to W (Figs. 2A-C). Global long-range synchrony was again observed to increase from W to N1 (Fig 2E, t=7.55, p<0.001) and to decrease from N1 to N2 (t=-9.16, p<0.001). Global long-range synchrony was not statistically different between W and N2 (t=-1.62, p=0.238). The relationship between mean regional local and long-range synchrony revealed low correlation within sleep states (Fig 2F; purple, green, and red; W, R^2^ = 0.204, p<0.001; N1, R^2^ = 0.109, p<0.001; N2, R^2^ = 0.098, p<0.001, respectively), which increased in correlation when comparing across states (Fig 2F; R^2^ = 0.158, p<0.001).

Within the alpha frequency band, long-range synchrony increased over the bilateral frontal and temporal regions with relative sparing of the parietal-occipital region in N1, as compared to W (Fig. 3s). The beta band demonstrated relatively increased synchrony in interhemispheric connectivity and anterior temporal regions, during N1 as compared to W (Fig. 3s). The frequency-specific activation patterns in N1 and N2 were similar; however, N1 overall demonstrated an increase in long-range neural synchrony as compared to N2 (Fig. 4s).

**Figure 3.**
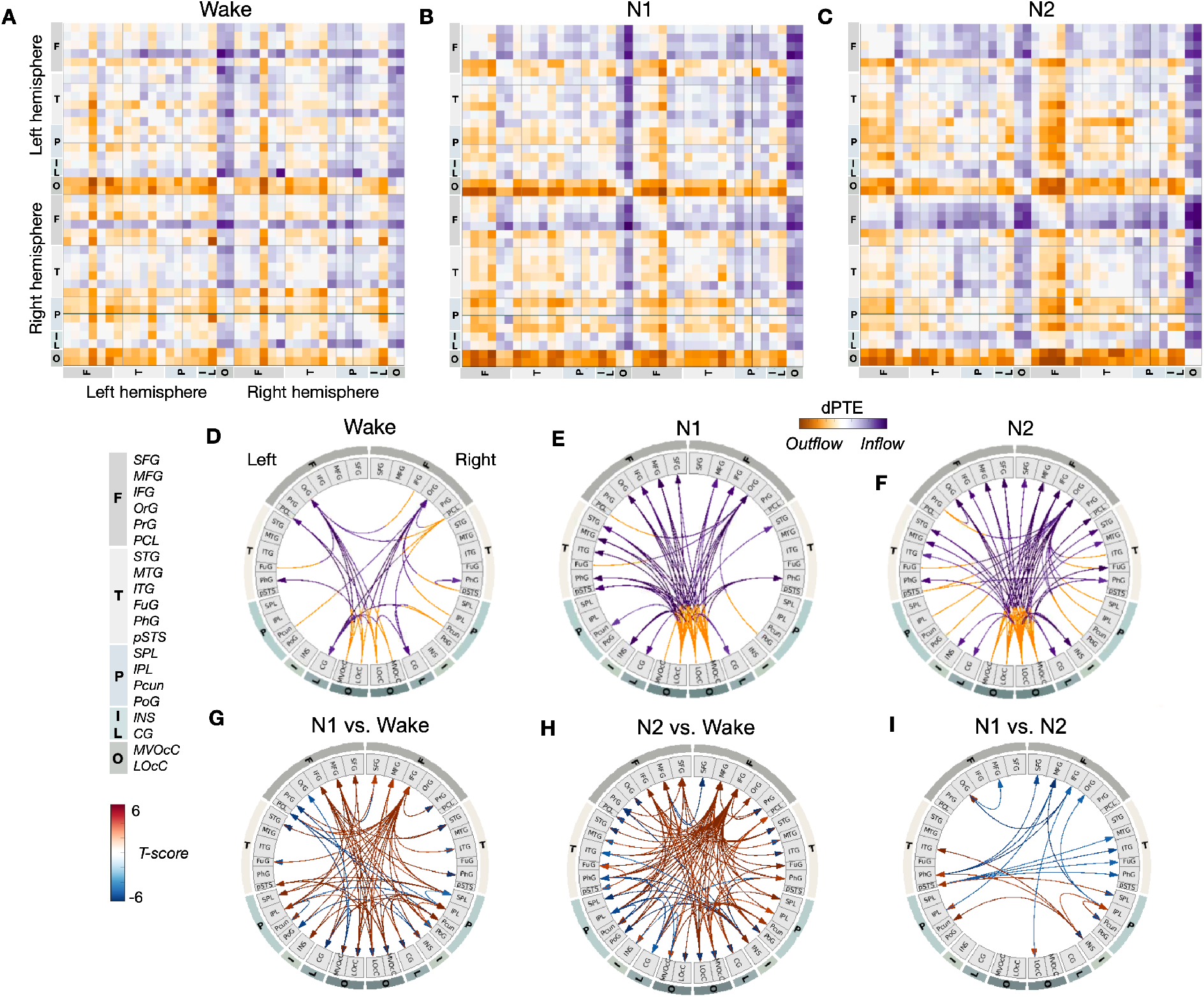
Directed information flow of sleep-wake states in the delta frequency band. A-C) Directed information flow as measured by directional phase transfer entropy (dPTE) across W, N1, and N2. From W to N1 and N2, there is increasing magnitude of dPTE with an aggregative inflow into the bilateral frontal regions. D-F) Alternate visualization of directed information flow colored by orange (outflow) and purple arrow (inflow). The visualization across W, N1, and N2 using a common threshold for depicting increased magnitudes and trajectories of the directed information flow. G-I) T-scores plotted indicating magnitude changes between specified states (accounting for FDR level 0.5)

**Figure 4.**
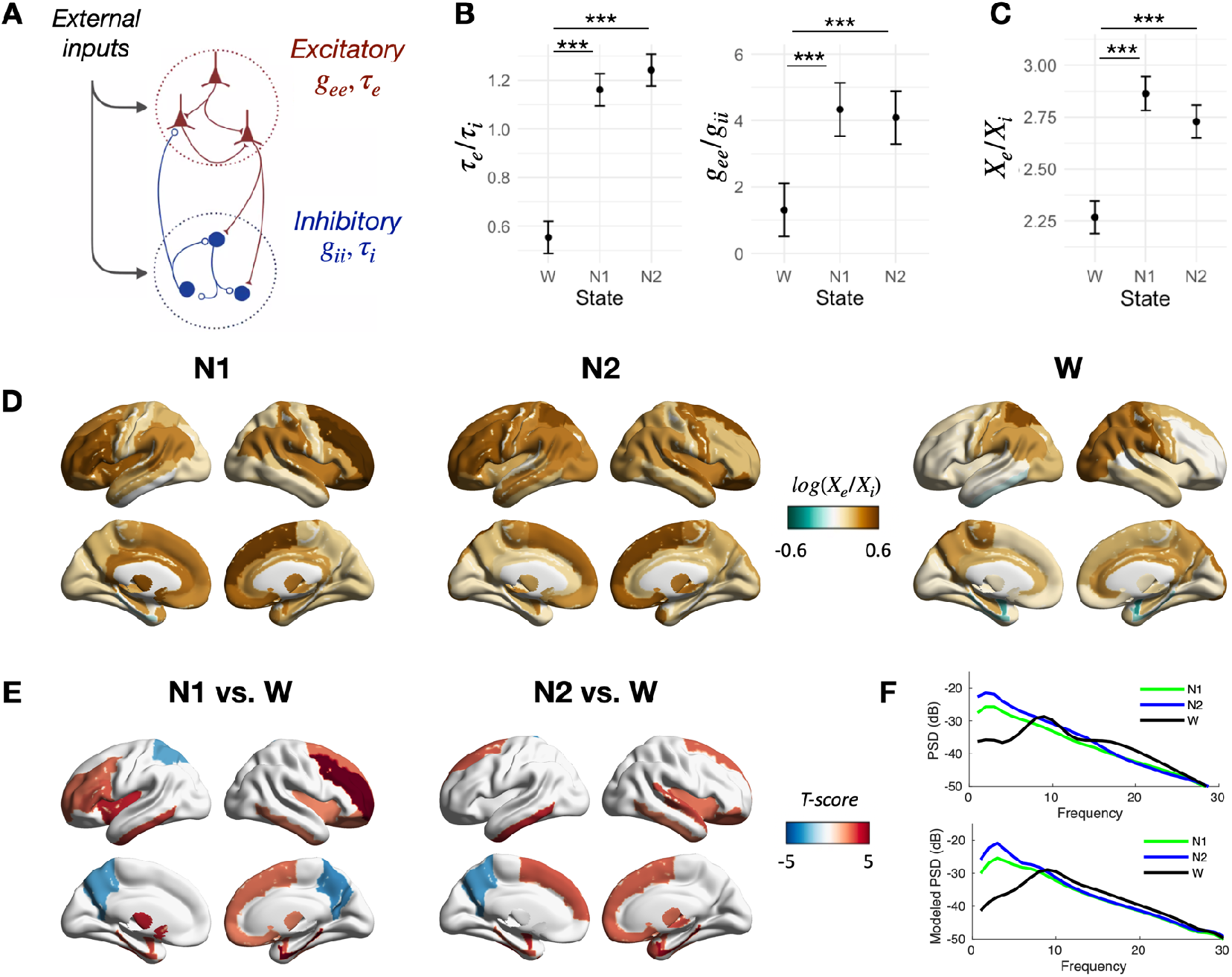
Biophysical NMM model across sleep-wake states demonstrates spatially distinct excitation-inhibition patterns with transition to sleep. A) Depiction of NMM, comprised of both excitatory and inhibitory inputs and modeled with time constants (*τ*) and gain (*g*) parameters. B) Optimized parameters of *τ_e_/τ_i_* and *g_ee_/g_ii_* across sleep-wake states. C) Excitatory-inhibitory ratios (*X_e_/X_i_*) across sleep-wake states, revealing an increased excitation-to-inhibition with transition from wake to sleep. D) Spatial maps of regional *X_e_/X_i_* across sleep-wake states, revealing cortical heterogeneity in excitation-to-inhibition. E) T-scores reflecting differences between N1 vs. W (*left*) and N2 vs. W (*right*) across different ROIs. Depicted T-scores are thresholded to account for multiple comparison testing (FDR level 0.05). F) Actual (*top*) and modeled (*bottom*) power spectral density curves across sleep-wake states averaged across all ROIs.

### Information flow across sleep-wake states

Information flow, as measured by dPTE^26^, provides insight on the regional inputs and outputs that may influence local activity and synchronization. Within the delta band, aggregated inflow tracts targeting bilateral frontal regions increased in magnitude when transitioning from W to N1 and from N1 to N2 (Fig. 3). Outflow from the bilateral occipital cortices (e.g. bilateral occipital polar cortex, lingual gyrus, inferior occipital gyrus, middle occipital gyrus, and ventromedial parietooccipital sulcus) were largely directed to the orbital frontal cortices during wakefulness and to bilateral frontal regions more broadly during N1 (Figs. 3B,E,G). Strikingly, with transition to N2, the outflow from the posterior regions gave rise to primary inputs not only to bilateral frontal regions, but also to the bilateral temporal lobe, insula, and cingulate, which then provided inflow into the bilateral frontal region (Fig. 3C,F,H). The serial flow of information, in which the bilateral frontal regions receive both direct and indirect inputs from broad cortical regions, represents a hierarchical organization of information flow. Given the spatial patterns during sleep, we speculate that the prominent anterior delta power seen at sleep onset may be enabled through the inputs of broadly distributed and hierarchically organized cortical activity.

In the alpha frequency band, there was an overall reduction in the occipital-frontal information flow (Fig. 5s) with transition from wake to NREM. In addition, the strong outflow tracks seen from the precuneus and IPL attenuated with the transition from W to N1 and from N1 to N2 (Fig. 5s). Importantly, these findings suggest that with the transition to sleep, the posterior-anterior information flow does not cease, but rather that there is a spectral shift to lower frequencies.

### Neural mass modeling of cortical excitation-to-inhibition parameters across sleep states

Finally, we utilized the observed physiology as inputs into biophysical NMMs to infer underlying mechanisms. Parameters of cortical excitation and inhibition were determined by modeling the observed regional spectral power using a deterministic NMM that captures local oscillatory dynamics (Fig 4A). The two indices of the NMM that were optimized to fit local electrophysiology included the characteristic time constants (*τ_e_, τ_i_*), representing the time-course of the underlying inhibitory and excitatory drive, and the gain (*g_ee_, g_i_*), representing the strength of recurrent local circuits. From wake to light NREM, both the *g_ee_/g_ii_* and the *τ_e_/τ_i_* ratio was observed to increase; differences between N1 to N2 were not statistically significant (Fig. 4B). The mean excitation-to-inhibition ratio (*X_e_/X_i_*), averaged across all regions, revealed an increase in excitation-to-inhibition from wake to light NREM (N1-W, t=10.4, p<0.001; N2-W, t=8.2, p<0.001) and a trend toward decreasing magnitudes from N1 to N2, although non-significant (N1-N2, t=2.3, p = 0.052; Fig 4C). These findings suggest that the transitional period to sleep may comprise a complex excitation-to-inhibition balance, in which *local* excitability-to-inhibition transiently increases with the transition to light NREM.

By modeling local dynamics, the excitation-to-inhibition ratio was furthermore demonstrated to be spatially distinct across the cortical surface within sleep-wake states (Fig 4D). The excitation-to-inhibition balance was highest over the bilateral frontal lobes during N1, while relatively reduced during wakefulness (Fig 4E). Furthermore, the bilateral parietal regions exhibited relatively increased cortical excitation-to-inhibition during wakefulness, as compared to light NREM (Fig. 4E). Comparison of the observed spectra to the model spectra (Fig. 4F) revealed an overall similarity in the profile of the spectra and peak frequencies. As homeostatic regulation during sleep has been thought be achieved through tuning synaptic strengths to alter the excitation-to-inhibition balance^27^, our computational modeling suggests that the sleep transition may support a spatially heterogeneous homeostatic cortical process.

## DISCUSSION

The characterization of whole-brain neural oscillations is essential to advancing our understanding of distributed network interactions underlying the transition to sleep. Descriptions of large-scale brain networks have relied chiefly on fMRI/PET imaging, which lack the temporal resolution and access to electrophysiology, and on EEG, which traditionally has limited spatial coverage and sensor level analysis. By using MEG imaging, a whole-brain, high-density imaging methodology, to simultaneously evaluate spectral power, functional connectivity, and information flow, we identify local and long-range cortical structure that underlie the transition from waking to light NREM sleep. Our findings suggest that the transition to sleep is accompanied by specific cortical patterns of neural synchrony that are coordinated in frequency and spatial extent. Furthermore, using directed measures of information flow, we identify a hierarchical flow of information from broad cortical areas to the bilateral frontal regions through both direct and indirect inputs. Finally, by employing a NMM, we demonstrate spatially heterogeneous excitatory-to-inhibitory parameters upon the transition to sleep, potentially suggestive of local homeostatic processes. As a simultaneous evaluation of local and long-range synchrony and information flow has not yet been performed during sleep onset, we review the literature in each of these domains in the context of our findings and then elaborate on how our NMM modeling fits in with experimental techniques.

Studies from the past decade have revealed that sleep onset is not a homogenous process^3,28–31^. Seminal PET studies provided the initial evidence of differential deactivation of cortical association regions with transition from waking to NREM, including in the prefrontal cortex, anterior cingulate, precuneus and mesial temporal structures, in addition to the deactivation of subcortical structures^32,33^. Slow wave activity using EEG/MEG has been identified to originate in anterior medial regions^28,34^, later involving parietal, temporal and occipital regions in an anterior to posterior direction^28^. The onset of slow wave activity coincides with an early rise in delta power in the frontal regions^21,23,34^ and an increase in theta synchrony in the occipital region^29,34^. These findings are similarly recapitulated in our data (Figs. 1B), in which the spectral reconstructions revealed prominent delta power in the mesial frontal regions. Across all frequency groups, regional local and long-range synchrony were poorly correlated (e.g. Figs. 1H, 2F), suggesting that the degree of connectivity has minimal bearing on the observed local synchrony. Similar findings in rest-state between relative power and mean PLI in specific bands have been previously identified^35^, reaffirming the need to simultaneously evaluate both local and long-range synchrony measures. In contrast, we did identify a strong correlation between regional local and long-range synchrony *across* sleep-wake states, suggesting that a global factor related to state-change impacts both local and long-range synchrony. A possible physiologic interpretation includes an increase in global mean spiking rates, elicited from subcortical inputs or achieved through a change in effective membrane time constants or integration time, leading to an increase in both local and long-range synchrony^36^.

Large-scale functional connectivity studies using fMRI^16,17,20^ and MEG^24^ have demonstrated widespread increases in long-range synchrony from W to N1 and N1 to N2, and a subsequent decrease in long-range synchrony with transition to N3^16,17,37^. The widespread synchronization has been associated with a decrease in global integration^38^, as measured by graph theoretic metrics, and theorized to account for loss of consciousness with transition to sleep. Although our findings also demonstrate an increase in mean long-range synchrony from W to light NREM, our findings further reveal distinct long-range interactions during light NREM, such as predominant bifrontal and fronto-parietal interactions in the delta band (Figs. 1D, E), and fronto-occipital and fronto-parietal interactions in the theta band (Figs. 2B, C). These findings differ from prior MEG studies^24^, which revealed homogenous increases in coherence, possibly due to limitations of sensor-based FC mappings, which may be confounded by overlapping lead fields and potentially conceal underlying structure. As compared to spectral power, the mean long-range synchrony of N1 was identified to be higher than both W and N2, consistent with prior literature^16,17,24,39^. These findings suggest that N1 composes a unique state that is not a transitional or ill-defined state between wake and sleep, which has previously been suggested due to the relatively low interrater reliability of identifying N1 based on less well-defined PSG features^40^ and challenges of machine learning fMRI studies to extract unique N1 states^41,42^. Rather, its distinctly elevated global long-range synchrony speaks to the complexity of the N1 state, which has been characterized by unique behavioral observations, including sleep mentation, responsiveness to sensory stimuli, or lucid dreaming^4,43^.

Complementing the evaluation of local and long-range synchrony across regions, we also assessed the directionality of regional interactions, as measured by information flow. Dominant posterior to anterior patterns of information flow of alpha frequencies have previously been identified during resting state^44^. Our findings confirm the posterior to anterior patterns of information flow during resting state for higher frequencies (Fig. 4s), and furthermore newly demonstrate increasing organization and magnitudes of information flow within the delta frequency with the transition from wake to light NREM. Specifically, information flow within the delta frequency composed of 1) a direct posterior to anterior pattern (i.e. from parietal-occipital regions to frontal regions), and 2) an indirect pattern (i.e. from posterior regions to bilateral temporal regions, which then flow to the bilateral frontal lobes; Figs. 3C, F), which together depict a hierarchical sequence of information flow during light NREM^44,45^. In contrast to prior EEG-based studies revealing an inversion of directionality from occipital-to-frontal to frontal-to-occipital during sleep onset^22,46,47^, our findings suggest a spectral shift in the prominent organizational pattern, while maintaining the posterior-to-anterior information flow. In addition to the increased number of sensors for source reconstruction approaches, MEG provides a reference free methodology for computing information flow, thereby preventing the potential bias associated with referenced based methods such as EEG.

Finally, we investigated whether the observed neural oscillatory patterns may give rise to spatially heterogeneous model parameters of cortical excitation-to-inhibition. Cortical excitability has been demonstrated to increase with time awake and sleep deprivation^48,49^ and decrease during sleep and the circadian cycle^49,50^, as measured by neuronal firing rates, synchronization, and propagation of transcranial magnetic stimulation (TMS)^49,51^. Our NMM modeling suggests that the transition from wake to light NREM is accompanied by a global increase in *local* cortical excitation-to-inhibition activity, consistent with prior methods using TMS^52^. Specifically, prior TMS findings revealed an enhancement of early or local TMS-EEG responses during sleep, followed by a global attenuation reflecting an overall global breakdown in effective cortical connectivity^52^. The amplified early or local TMS-EEG response has been hypothesized to relate to an increased drive of post-synaptic excitatory neurons, synchronization of local cortical populations or activation of thalamocortical circuits^52^. While our model was assessed in a hypothesis-generating framework, further studies are required for validation of this model. Additionally, the computational model used here only accounts for local synchrony patterns; methods to incorporate both local and long-range connectivity within a parsimonious model are currently being developed and would be of future interest to incorporate given the distinct local and long-range corticocortical synchrony patterns observed.

By employing local NMM modeling and leveraging whole-brain local field potential, cortical spatial maps of excitation-to-inhibition parameters can be elucidated, in contrast to seeded processes, e.g. TMS impulses^49,51,53^, magnetic resonance spectroscopy^54^, or direct measurements of excitatory and inhibitory neurotransmitters^55^. As such, we demonstrate that the modeled ratio of excitation-to-inhibition across the cortex is non-uniform (Fig. 4D, E) and is highest over the bilateral frontal lobes during light NREM. In contrast, the highest excitation-to-inhibition ratio during wakefulness is within the parietal-occipital regions. Our findings support the view that sleep is composed of spatially heterogeneous network dynamics, which may reflect complex homeostatic patterns across the cortex.

This work has limitations. Due to the noisy environment and time-limitations of a recording session within the MEG scanner, the sleep quality in the healthy participants was at times fragmented. In addition, deep NREM stages were not obtained across all patients and thus not included in the analysis. To address the sleep fragmentation, shortened epochs of high-quality 15s were utilized and concatenated together to formulate the most faithful representations of sleep-wake states. Future efforts may benefit from overnight or sleep-deprived recording sessions to obtain longer, continuous periods of sleep-wake states. This study also incorporates a modest sample size of fourteen patients, who are also older in age. A future research direction includes the characterization of differences in neural synchrony patterns during sleep across age. In addition, a fundamental limitation of current human electrophysiology recordings is the lack of simultaneous recordings from both whole-coverage cortical areas and subcortical structures, including the thalamus. Thus, the contributions of subcortical structures to the cortical electrophysiologic synchrony patterns observed here remains an open question. Finally, although the linear NMM captures the frequency spectra directly through a closed form solution, the primary limitations of the NMM employed in this study include its inability to capture non-linear complexities of the brain.

In summary, our work identifies cortical oscillations structured across multiple spatial scales and hierarchically organized across time (e.g. frequency-specific interactions) with the transition to light NREM sleep. Serial information flow from broad cortical regions are directed to the mesial frontal regions in the delta frequency, which may contribute to the synchronization of local oscillations. Finally, our observed electrophysiology gives rise to spatially heterogeneous parameters of cortical excitation-to-inhibition with transition to sleep. Together, our findings suggest that the transition to sleep is accompanied by distinct, whole-brain, frequency and spatially-specific neural patterns that underlie spatially heterogeneous excitatory and inhibitory activity.

## METHODS

### Study cohort

MEG and simultaneously obtained scalp EEG data were obtained from fourteen healthy elderly individuals (age, mean 63.6 y [SD 11.2]; gender, female 11 of 19 [78.6%]). All participants underwent a complete history and physical examination. Exclusion criteria for this healthy cohort included any neurologic disorders by history or abnormal neurologic findings by exam. In addition, exclusion criteria included use of any chronic or acute neuroactive medications, including neuroleptics, benzodiazepines, SSRI/SNRIs, or sedatives. Inclusion criteria included achieving at least an aggregate of 60 seconds within each sleep-wake state (W, N1 and N2, segmented into 15 second epochs, see below) during the EEG/MEG study. In all subjects, the use of these data for research was approved by the University of California Institutional Review Board, and all subjects provided written informed consent prior to data collection.

### Data acquisition and preprocessing

EEG/MEG data were obtained for a period of 50-70 minutes of resting state, recorded at a sampling rate of 600 Hz. Before the start of the recording, subjects were instructed to close their eyes and attempt to sleep. Subjects did not engage in any tasks during the recording. Raw EEG/MEG traces were chunked into 15 second epochs. Each epoch was visually inspected and rejected if there was presence of movement artifacts. In a subset of subjects with metallic artifacts from non-cranial implants, a digital signal subspace project (DSSP) algorithm was used for denoising^56^. Epochs were scored into the appropriate sleep states based on scalp EEG criteria and using two-person validation according to American Academy of Sleep Medicine (AASM) criteria. The 15 second epochs of MEG data were then concatenated together to achieve at minimum, a 60 second time series representing each behavioral state. A maximum of the first eight artifact-free epochs, e.g. 120 second sensor time, were utilized in this analysis. Prior work demonstrated that 60 seconds of resting wake state is an adequate duration to reliably achieve stationarity for MEG time series^57^. Source reconstruction was performed using adaptive spatial filtering methods^58^ at an individual subject level, yielding voxel level source time series. Source reconstructions were co-registered to MNI template, and regional level time courses were extracted with alignment to the Brainnetome atlas^59^, parcellated into 210 cortical regions of interests (ROIs). The data analysis pipeline is illustrated in Fig 1s.

### Data analysis and neural mass model

Time series data were bandpass filtered into the following frequency bands: delta (1-4 Hz), theta (4-8 Hz), alpha (8-12 Hz), sigma (12-15 Hz), and beta (15-30 Hz). Local neural synchrony was quantified by the normalized spectral power of the canonical frequency bands within a given ROI. Normalized spectral power was determined by dividing the power spectra by the total power.

Long-range synchrony was computed using imaginary coherence, an established spectral coherence measure of neural synchrony that is robust to volume conduction effects^60^. Imaginary coherence was computed across pairwise atlas-based regions to assess for long-range synchrony patterns; group statistics are described below. Mean regional long-range synchrony was computed by averaging the pairwise long-range synchrony measures for an individual region, i.e. averaging the pairwise imaginary coherence matrix along one axis. Global long-range synchrony was computed as the average of all regional long-range synchrony. To visualize regional patterns of imaginary coherence, spatial maps of the magnitudes of regional long-range synchrony were visualized on the Brainnetome atlas. We note that imaginary coherence is a shrinkage estimator of true synchrony that includes both real and imaginary components of coherence. Although near-zero lag can be observed in large scale oscillatory networks^61,62^, imaginary coherence is favored given the concern for additional non-biological synchrony that may be captured in zero-lag synchrony.

To quantify information flow, phase transfer entropy (PTE) was also computed to assess the pairwise directional interaction between ROI time courses^26^, and directional information was computed using directional phase transfer entropy (dPTE), a connectivity measure for quantifying information flow that is robust to noise and linear signal mixing^63^. Data analysis was performed using the FieldTrip Matlab Toolbox^64^ and custom made MATLAB code (version R2019a).

In addition, we utilized a linear, deterministic neural mass model (NMM)^65–68^ to gain mechanistic insights on the relative balance of excitatory and inhibitory inputs based on the observed spectra. This NMM captures the local neural activity, as measured by the power spectral density (PSD) across each ROI and sleep-wake state. In this implementation of the NMM, whole-brain dynamics are deterministically modeled in closed form within the Fourier domain, using a canonical rate model that considers local cortical neural activity^65,66^. NMM methods are further described in Ranasinghe et al^69^. Specifically, we modeled a local signal as the summation of excitatory signals *x_e_*(*t*) and inhibitory signals *x_i_*(*t*) for every ROI, based on the Brainnetome parcellation^59^. Excitatory and inhibitory signals were modeled by a decay of the respective signals, feedback inputs from both excitatory and inhibitory signals, and noise. Parameters for the excitatory and inhibitory signals for individual regions included the neural gains for the excitatory, inhibitory, and alternating populations represented by *g_ee_*, *g_ii_*, and *g_ei_*, respectively; the characteristic time constants of excitatory and inhibitory populations represented by *τ_e_* and *τ_i_*, respectively; the Gaussian white noise term represented by *p*(*t*); and the Gamma-shaped ensemble neural impulse response, represented by *f_e_*(*t*) and *f_i_*(*t*) for inhibitory and excitatory contributions, respectively. The dynamics of the excitatory and the inhibitory signals are thus represented by the following:

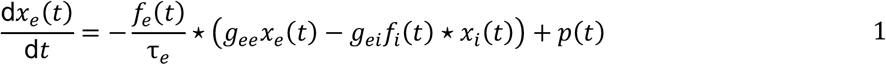

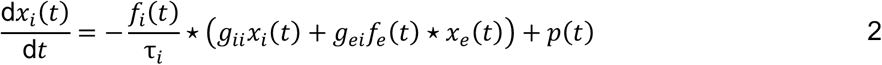

For each ROI of the Brainnetome atlas, *g_ee_, g_ii_*, *τ_e_*, and *τ_i_* were estimated, while *g_ei_* was fixed at 1. Optimization of the model was based on mean squared errors (MSE) between the actual and modeled PSD curve, for each ROI, participant, and sleep-wake state. The closed-form expression of the modeled PSD is described further in the Supplementary Methods. The minimization of the MSE for parameter optimization was performed using the basin hopping global optimization algorithm (Python)^70^. An empirical cumulative distribution function was constructed on the basis of MSE across all regions of interest, sleep-wake states, and patients; outliers were determined based on the threshold inclusion of 75% of the cumulative distribution (corresponding to MSE 37.6). Subsequent statistical testing to compare model parameters across sleep-wake states was performed using a repeated mixed models approach on a modular ROI level, accounting for subject, ROI, frequency, and state.

### Group statistics

For statistical comparisons across states, the 210 cortical regional parcellation level spatial maps of the Brainnetome atlas were configured to 44 modular level parcellation maps, giving rise to 990 functional connections, which includes the averaged measure (e.g. imaginary coherence) within each module. Linear mixed effects models with repeated measures were performed to identify statistically significant differences across two states for modules and modular connections. Statistics were corrected for multiple comparisons (false discovery rate, FDR, level 0.05).

## ACKNOWLEDGEMENTS

We thank our subjects for volunteering to participate in the study.

## DATA AVAILBABILITY

Anonymized summary data and relevant code will be made available upon reasonable request.

## AUTHOR CONTRIBUTIONS

J.M.F., K.K., P.V., K.G.R., H.E.K., A.R., A.D.K., and S.S.N. contributed to the conception of the work, analysis, and interpretation of the data. H.M. contributed to the analysis of the data. A.M.F., K.G.R., and K.V. contributed to the acquisition of the data. J.M.F., P.V., A.D.K., S.S.N. contributed to the writing of the manuscript. All authors have reviewed and approved the submitted manuscript.

## Abbreviations

Amy: amygdala
CG: cingulate gyrus
FC: functional connectivity
FuG: fusiform gyrus
Hipp: hippocampus
IFG: inferior frontal gyrus
IPL: inferior parietal lobule
INS: insula
ITG: inferior temporal gyrus
IQR: interquartile range
LocC: lateral occipital cortex
MEG: magnetoencephalography
MFG: middle frontal gyrus
MTG: middle temporal gyrus
MVOcC: medioventral occipital cortex
NREM: non-rapid eye movement
N1: stage 1 sleep (NREM)
N2: stage 2 sleep (NREM)
OrG: orbital frontal gyrus
PCL: paracentral lobule
Pcun: precuneus
PhG: parahippocampal gyrus
PoG: postcentral gyrus
PrG: precentral gyrus
pSTS: posterior superior temporal sulcus
ROI: region of interest
SFG: superior frontal gyrus
SPL: superior parietal lobule
STG: superior temporal gyrus
W: wake

