## Supplementary Methods and Materials for "Cortical structure of neural synchrony and information flow during transition from wakefulness to light non-rapid eye movement sleep"

### SUPPLEMENTARY METHODS AND FIGURES

#### Methods

1. Neural mass modeling expanded methods, including derivation of closed form expression of modeled PSD

#### Figures

1. Methods flow chart
2. Spatial map of all spectral changes across state
3. Long-range synchrony changes for alpha/beta frequencies
4. Long-range synchrony changes between N1 and N2
5. Information flow for alpha frequency

### Supplementary Methods

#### Complete mathematical modeling and parameter estimation description

We used a linear neural mass model (NMM)[67,68] based on prior work from our group[65,66] and recapitulated in Ranasinghe et al[69] to gain mechanistic insights on the relative balance of excitatory and inhibitory signals based on the observed spectra. As in the main text, we modeled a local signal as the summation of excitatory signals $x_e(t)$ and inhibitory signals $x_i(t)$ for every ROI, based on the Brainnetome parcellation[59]. Excitatory and inhibitory signals were modeled by a decay of the respective signals, feedback inputs from both excitatory and inhibitory signals and noise. Parameters for the excitatory and inhibitory signals for individual regions included the neural gains for the excitatory, inhibitory, and alternating populations represented by $g_{ee}$, $g_{ii}$, and $g_{ei}$, respectively; and the characteristic time constants of excitatory and inhibitory populations represented by $\tau_e$ and $\tau_i$, respectively. This model also included the Gaussian white noise term represented by $p(t)$; and the Gamma-shaped ensemble neural impulse response, represented by $f_e(t)$ and $f_i(t)$ for inhibitory and excitatory contributions, respectively. The excitatory and the inhibitory signals are thus represented by

$$\frac{\mathrm{d}x_e(t)}{\mathrm{d}t} = -\frac{f_e(t)}{\tau_e} \star \left(g_{ee}x_e(t) - g_{ei}f_i(t) \star x_i(t)\right) + p(t) \qquad 1$$

$$\frac{\mathrm{d}x_i(t)}{\mathrm{d}t} = -\frac{f_i(t)}{\tau_i} \star \left(g_{ii}x_i(t) + g_{ei}f_e(t) \star x_e(t)\right) + p(t) \qquad 2$$

where $\star$ stands for convolution and the excitatory and inhibitory neural impulse functions are given as:

$$f_e(t) = \frac{t}{\tau_e^2} e^{\frac{-t}{\tau_e}} \qquad 3$$

$$f_i(t) = \frac{t}{\tau_i^2} e^{\frac{-t}{\tau_i}} \qquad 4$$

By taking the Fourier transform of Equations 1 and 2, $x_e(t)$ and $x_i(t)$ are then transformed into the Fourier domain and are expressed by $X_e(\omega)$ and $X_i(\omega)$, respectively, in which ω is the frequency:

$$j\omega X_e(\omega) = -\frac{F_e(\omega)}{\tau_e}\left(g_{ee}X_e(\omega) - g_{ei}F_i(\omega)X_i(\omega)\right) + P(\omega) \quad 5$$

$$j\omega X_i(\omega) = -\frac{F_i(\omega)}{\tau_i}\left(g_{ii}X_i(\omega) + g_{ei}F_e(\omega)X_e(\omega)\right) + P(\omega) \quad 6$$

where $j$ is the imaginary unit. Similarly, $P(\omega)$ represents the Fourier transform of $p(t)$, and $F_e(\omega)$ and $F_i(\omega)$ represent the Fourier transform of $f_e(t)$ and $f_i(t)$, and are expressed by:

$$F_e(\omega) = \frac{\frac{1}{\tau_e^2}}{\left(j\omega + \frac{1}{\tau_e}\right)^2} \quad 7$$

$$F_i(\omega) = \frac{\frac{1}{\tau_i^2}}{\left(j\omega + \frac{1}{\tau_i}\right)^2} \quad 8$$

Thus, the closed form solution of $X_e(\omega)$ and $X_i(\omega)$ is:

$$X_e(\omega) = \frac{\left(1 + \frac{\frac{g_{ei}}{\tau_e}F_e(\omega)F_i(\omega)}{j\omega + \frac{g_{ii}}{\tau_i}F_i(\omega)}\right)P(\omega)}{j\omega + \frac{g_{ee}}{\tau_e}F_e(\omega) + \frac{\left(g_{ei}F_e(\omega)F_i(\omega)\right)^2}{\tau_e\tau_i\left(j\omega + \frac{g_{ii}}{\tau_i}F_i(\omega)\right)}} \quad 9$$

$$X_i(\omega) = \frac{\left(1 - \frac{\frac{g_{ei}}{\tau_i}F_e(\omega)F_i(\omega)}{j\omega + \frac{g_{ee}}{\tau_e}F_e(\omega)}\right)P(\omega)}{j\omega + \frac{g_{ii}}{\tau_i}F_i(\omega) + \frac{\left(g_{ei}F_e(\omega)F_i(\omega)\right)^2}{\tau_e\tau_i\left(j\omega + \frac{g_{ee}}{\tau_e}F_e(\omega)\right)}} \quad 10$$

The simulated spectra, $X(\omega)$, is comprised of the excitatory and inhibitory components summed together, e.g. $X_e(\omega) + X_i(\omega)$. Using the transfer functions, $H_e(\omega)$ and $H_i(\omega)$, and $P(\omega)$ as the driving function, $X_e(\omega)$ and $X_i(\omega)$ can be re-expressed as $H_e(\omega)P(\omega)$ and $H_i(\omega)P(\omega)$, and thus, $X(\omega) = (H_e(\omega) + H_i(\omega))P(\omega)$. From here, the PSD is represented by $\mathbb{E}(|X(\omega)|^2)$. Recall that $P(\omega)$ represents Gaussian noise and thus has a flat power spectrum. Therefore, $\mathbb{E}(|X(\omega)|^2) \propto |H_e(\omega) + H_i(\omega)|^2$, which is subsequently converted to dB scale by log-transformation: $10\log_{10}(|H_e(\omega) + H_i(\omega)|^2)$.

For each ROI of the Brainnetome atlas, $g_{ee}$, $g_{ii}$, $\tau_e$, and $\tau_i$ were estimated, while $g_{ei}$ was fixed at 1. The spectra was modeled across frequencies 1-30 Hz, and the fit of the model was determined by calculating the mean squared error (MSE) between the simulated model PSD and the observed, source-localized MEG PSD in dB (with MEG dB spectra scaled by 2.5 to match the modeled spectra magnitude) across all frequencies. The minimization of the MSE for parameter optimization was then performed using the basin hopping global optimization algorithm (Python)[70]. The model parameter initialization value, upper-boundary, and lower-boundary were specified as 17 ms, 5ms, and 30 ms, respectively, for $\tau_e$ and $\tau_i$; and 5, 0.1 and 10, respectively, for $g_{ee}$ and $g_{ii}$. In addition, hyperparameters, including the number of iterations, step-size, and temperature, were 2000, 4, and 0.1, respectively. If boundary-values were hit during the parameter optimization, the step-size was augmented to 6 for that specific ROI. The parameters leading to the lower MSE was then selected. This optimization procedure was performed for each Brainnetome ROI, for each subject and sleep-wake state.

### Supplementary Figures

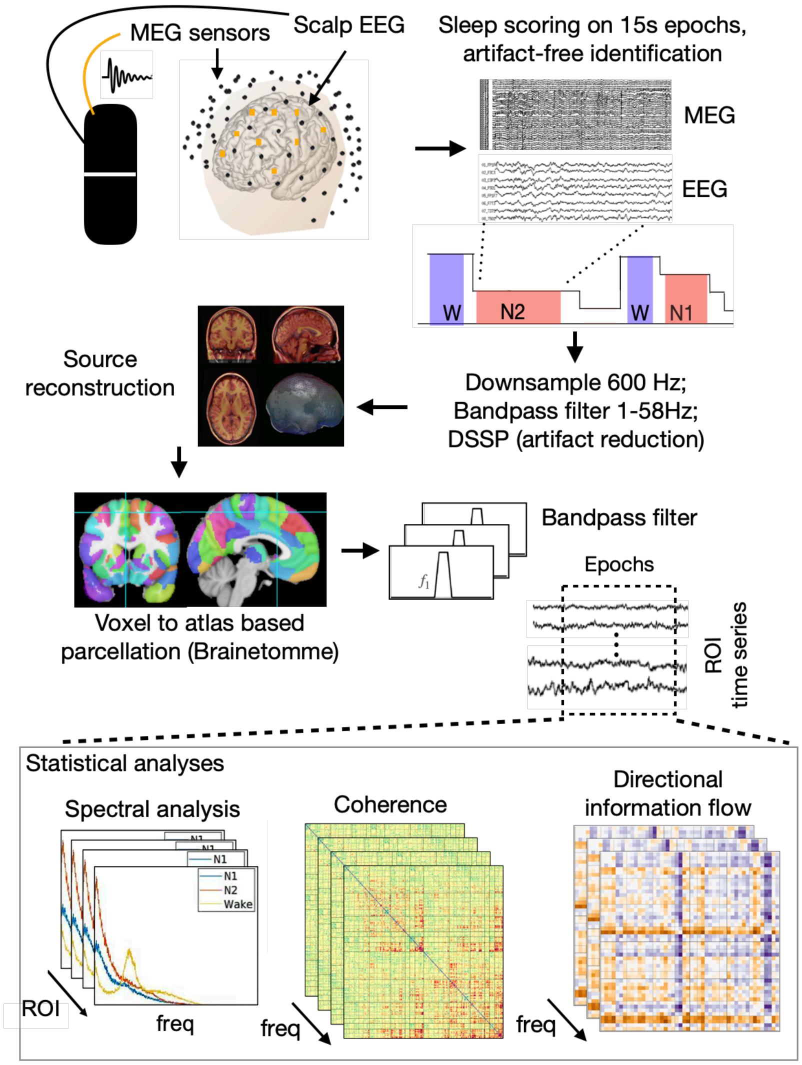


**Figure 1s. Methodology flow chart.** Neural signals are simultaneously acquired using MEG and scalp EEG sensors. The scalp EEG is chunked into 15s epochs, which is visually scored into sleep-wake states and reviewed for artifact. Time series (60s concatenated epochs) per state then undergo standard preprocessing steps and source reconstruction using adaptive spatial filtering methods[58] at an individual subject level, yielding voxel level source time series. The voxel level source time is mapped to the Brainnetome atlas. Bandpass filtering of the neural signal is performed on canonical frequency bands for further statistical analysis.

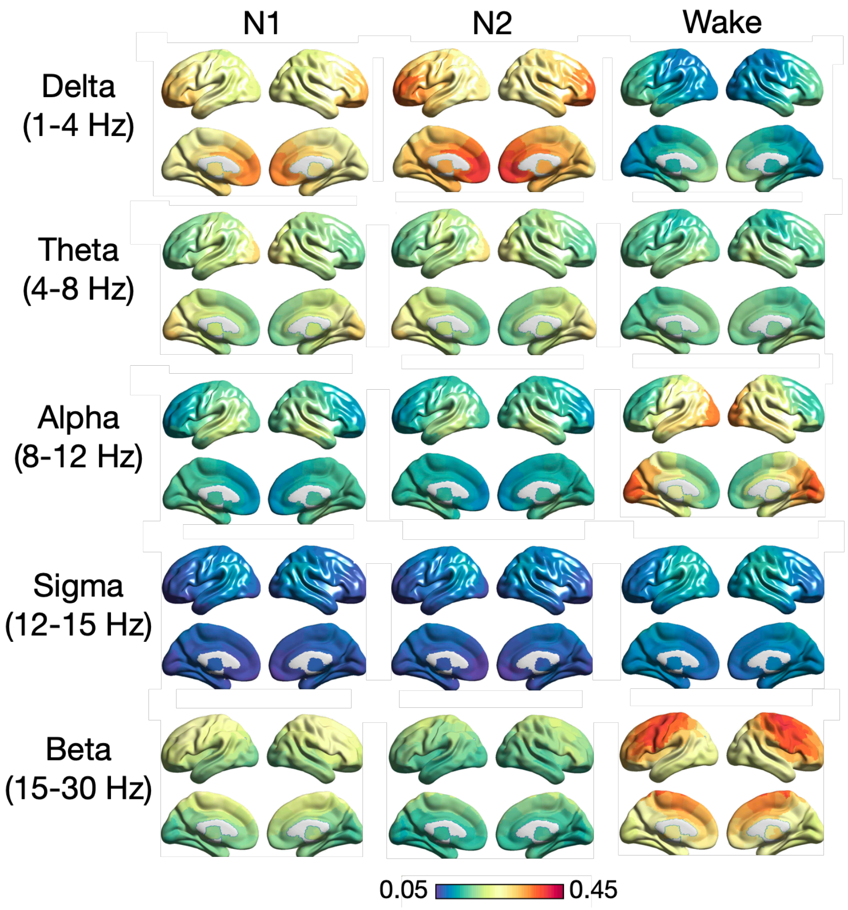


**Figure 2s. Spatial maps of normalized spectral power across sleep-wake states**. Spectral power is highest within the delta frequency band within N1 and N2, most prominent over mesial frontal regions. During wakefulness, there is an expected anterior-posterior distribution within the alpha frequency, most prominent within the bilateral occipital regions. In addition, there is strong spectral power within the beta band over the bilateral fronto-parietal regions during wakefulness.

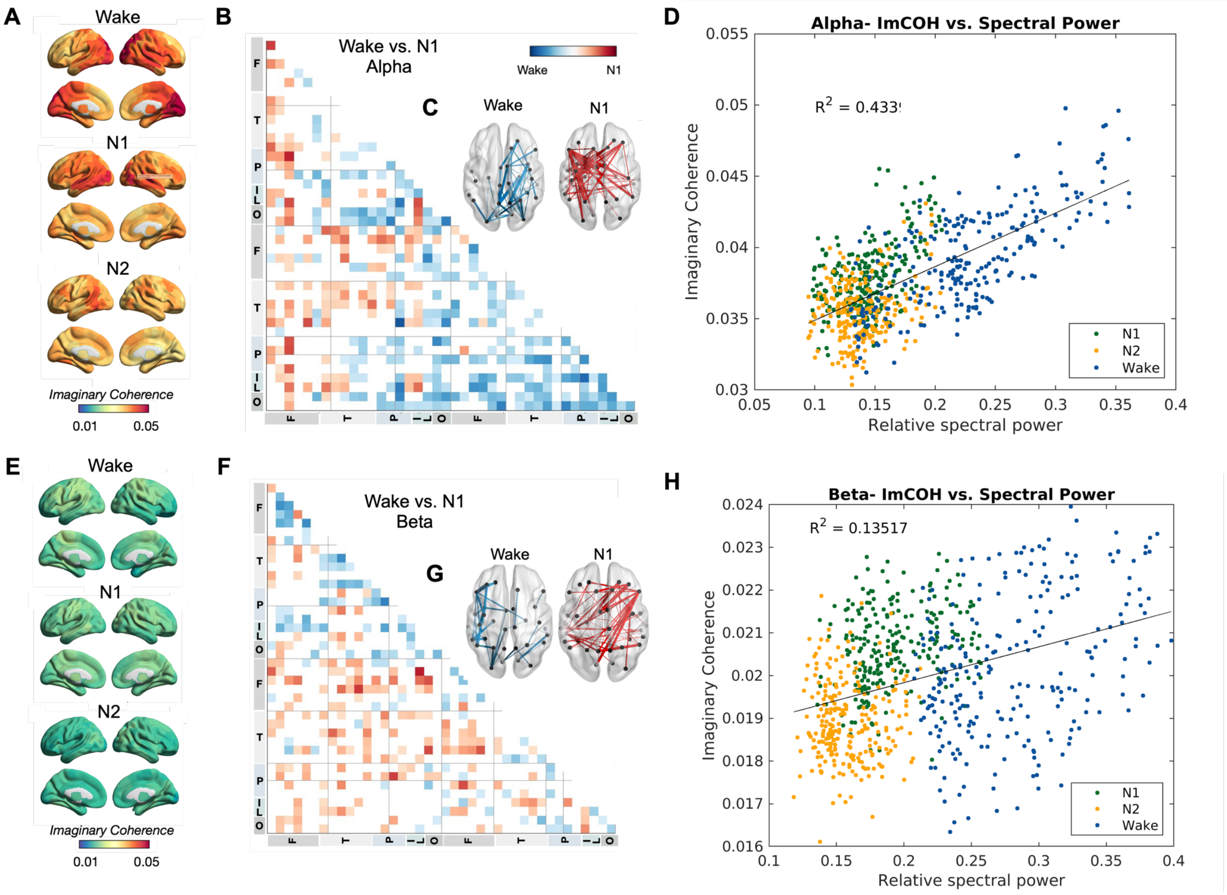


**Figure 3s. Spatial maps of mean long-range synchrony between sleep-wake states in alpha and beta frequency bands.** A) Spatial map of mean long-range synchrony in W, N1, and N2 within the alpha band, as measured by imaginary coherence. B) T-score map of differences in long-range synchrony between W and N1 within the alpha band. (C) Top 50 highest positive and negative functional connections, favoring N1 (red) and W (blue), respectively. D) Association between the mean regional local and long-range synchrony in the alpha band. E-H) Same as A-D, but for the beta frequency band.

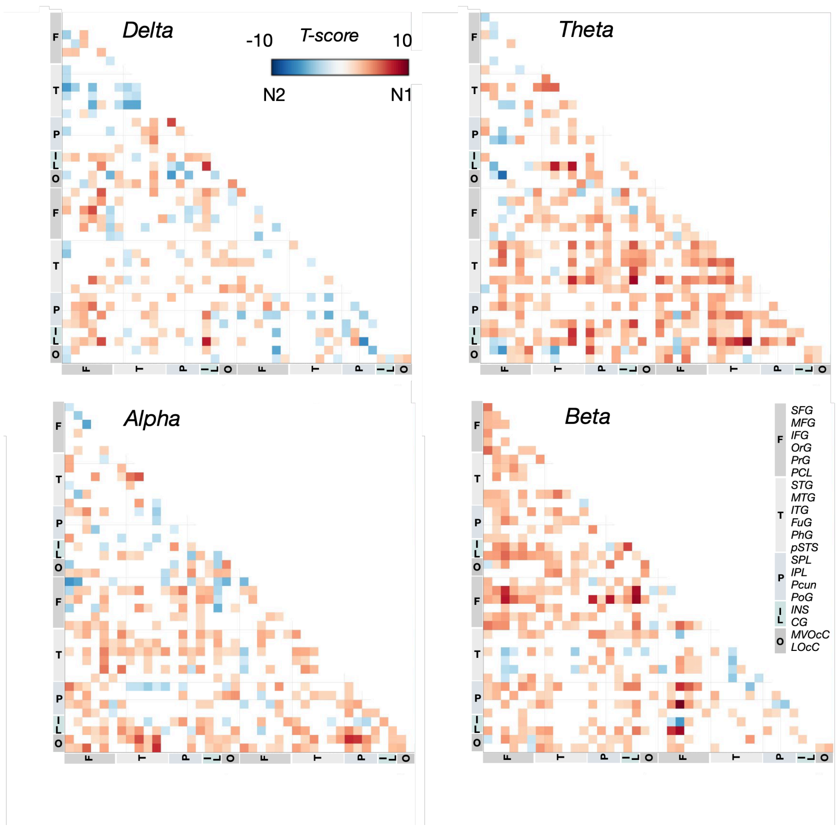


**Figure 4s. Spatial long-range synchrony maps N1 and N2.** T-score map of differences in long-range synchrony between N1 and N2 within the delta, theta, alpha and beta frequency bands. There is an overall preponderance of higher long-range synchrony in N1 (red), as compared to N2, across frequency bands.

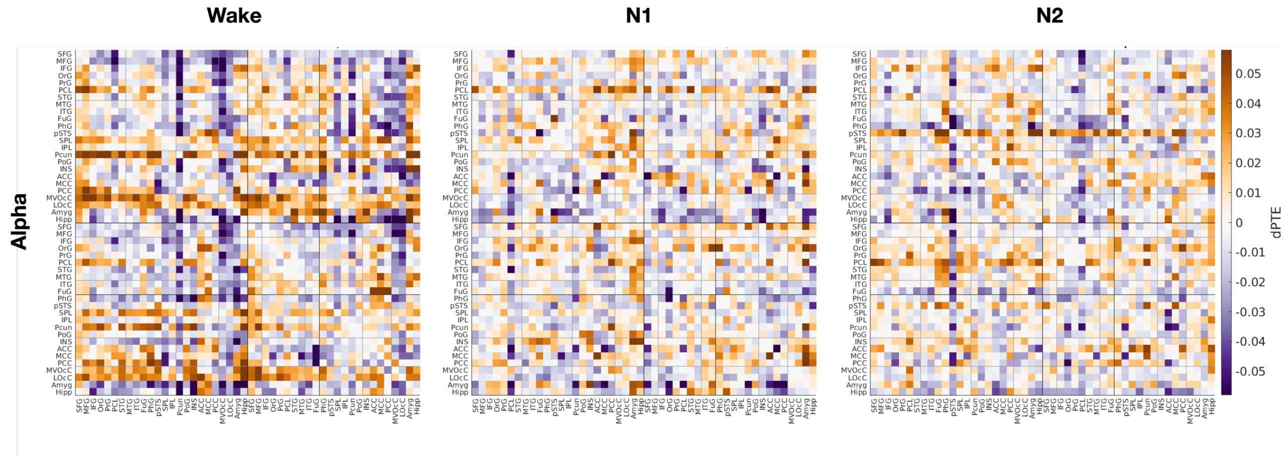


**Figure 5s. Information flow within the alpha frequency band**. Directed information flow as measured by directional phase transfer entropy (dPTE) across W, N1, and N2 for the alpha frequency band. From W to N1 and N2, there is an overall decrease in the magnitude of dPTE. Specifically, there is a reduction in outflow to inflow from the bilateral occipital regions (MVOcC, LOcC) and Pcun to the bilateral frontal regions (SFG, MFG) when transitioning from W to N1 and N2.
